# Behavioural evidence of spectral opponent processing in the visual system of stomatopod crustaceans

**DOI:** 10.1101/2024.05.01.592115

**Authors:** Ching-Wen Judy Wang, Justin Marshall

**Author notes:** Corresponding author’s. Competing interests: The authors declare no competing or financial interests.

## Abstract

Stomatopods, commonly known as mantis shrimps, possess an intricate colour vision with up to 12 photoreceptor classes organised in four specialised ommatidia rows (rows 1-4 in the midband region of the eye) for colour perception. While 2-4 spectral sensitivities suffice for most visual systems, the mechanism behind stomatopods’ 12-channel colour vision remains unclear. Based on neuroarchitecture, it was initially suggested that rows 1-4 may function as four parallel dichromatic channels allowing fine spectral discrimination and strong colour constancy in narrow spectral zones. Subsequently, unexpectedly low resolution in behavioural experiments indicated that a binning processing system may operate instead of or in addition to the ‘normal’ opponent processing system, categorising information into separate channels to create an activation pattern for rapid colour recognition. Previous anatomical and behavioural studies have speculated on the potential coexistence of these two systems in stomatopods’ colour vision. However, no behavioural study has specifically investigated the potential for colour opponency in their colour vision. Our findings provide the first direct behavioural evidence for spectral opponency in stomatopods’ visual system, showing that rows 1-4 operate, at least some of the time, as multiple dichromatic channels.

## Introduction

Stomatopods, also known as mantis shrimps, are marine crustacean predators inhabiting coral reef waters in tropical and subtropical environments. Having diverged from the other crustaceans approximately 400 million years ago, stomatopods have undergone an evolutionary path, separate to other crustaceans with notably the development of a sophisticated visual system (Manning, 1977; Marshall, 1988; Cronin and Marshall, 1989a; Cronin and Marshall, 1989b; Marshall et al., 2007; Marshall and Arikawa, 2014). Their visual system possesses the highest diversity of photoreceptors among all animals, with perhaps butterflies as close rivals (Arikawa et al., 1987; Koshitaka et al., 2008; Arikawa and Stavenga, 2014; Marshall and Arikawa, 2014), incorporating up to 20 types of photoreceptors used for various tasks including colour, polarisation and brightness detection. Their colour vision, is unique in particular, utilising as many as 12 classes of photoreceptors to sample chromatic information from deep UV to far red (300-750 nm) (Marshall, 1988; Cronin and Marshall, 1989b; Thoen et al., 2017b). As most animals use between 2-4 spectral sensitivities for colour vision, with more than 4 adding little discrimination ability (Barlow, 1982; Maloney, 1986), the reason for the apparent chromatic-complexity of the stomatopod system remains unclear.

The compound eye of most shallow water stomatopods comprises dorsal and ventral hemispherical regions, separated 6 rows of specialised ommatidia, known as the midband (Fig. 1A) (Marshall, 1988; Cronin and Marshall, 1989a; Porter et al., 2010). The dorsal four rows (rows 1-4) are dedicated to colour vision, while the last two rows (rows 5 and 6) serve for both circular (or elliptical) and linear polarisation vision in human-visible (around 500nm) and UV wavelengths, respectively (Fig. 1A, C) (Cronin and Marshall, 1989a; Cronin and Marshall, 1989b; Chiou et al., 2008; Thoen et al., 2017b). In addition to the midband, the hemispherical parts are employed for detecting human-visible linear polarisation and luminance information, exhibiting structures and functions similar to those in other crustaceans (Fig. 1A, C) (Cronin and Marshall, 1989a; Marshall et al., 1991; Kleinlogel et al., 2003; How and Marshall, 2014; Daly et al., 2016). In the stomatopod eye, each ommatidium consists of eight retinular cells, termed R1-R8. The architecture of ommatidia varies depending on their function (Marshall, 1988; Cronin and Marshall, 1989b; Marshall et al., 1991; Marshall and Oberwinkler, 1999; Marshall et al., 2007). Ommatidia responsible for polarisation vision demonstrate simpler structures, featuring two tiers, with a short R8 located distally to the main rhabdom R1-7 (Fig. 1C). In contrast, the ommatidia designated for colour vision are divided into three tiers, including R8 and two separate tiers of R1-7 (R1, 4, and 5 and R2, 3, 6 and 7). Within each ommatidium, every tier functions as a long-pass filter for the receptors underneath it, progressively narrowing the spectral sensitivities of these receptors. In Row 2 and 3, the filtering effect is further facilitated by intrarhabdomal filters positioned between the two tiers of R1-7. These filters consist of photostatic pigments that enhance the sharpness of spectral sensitivities, fine-tuning perceptions towards even longer wavelengths. As a result, each tier possesses a distinct spectral sensitivity, rendering the dorsal four rows of the midband an array of spectral channels, housing 12 classes of photoreceptors (Fig. 1B-D).

**Figure 1.**
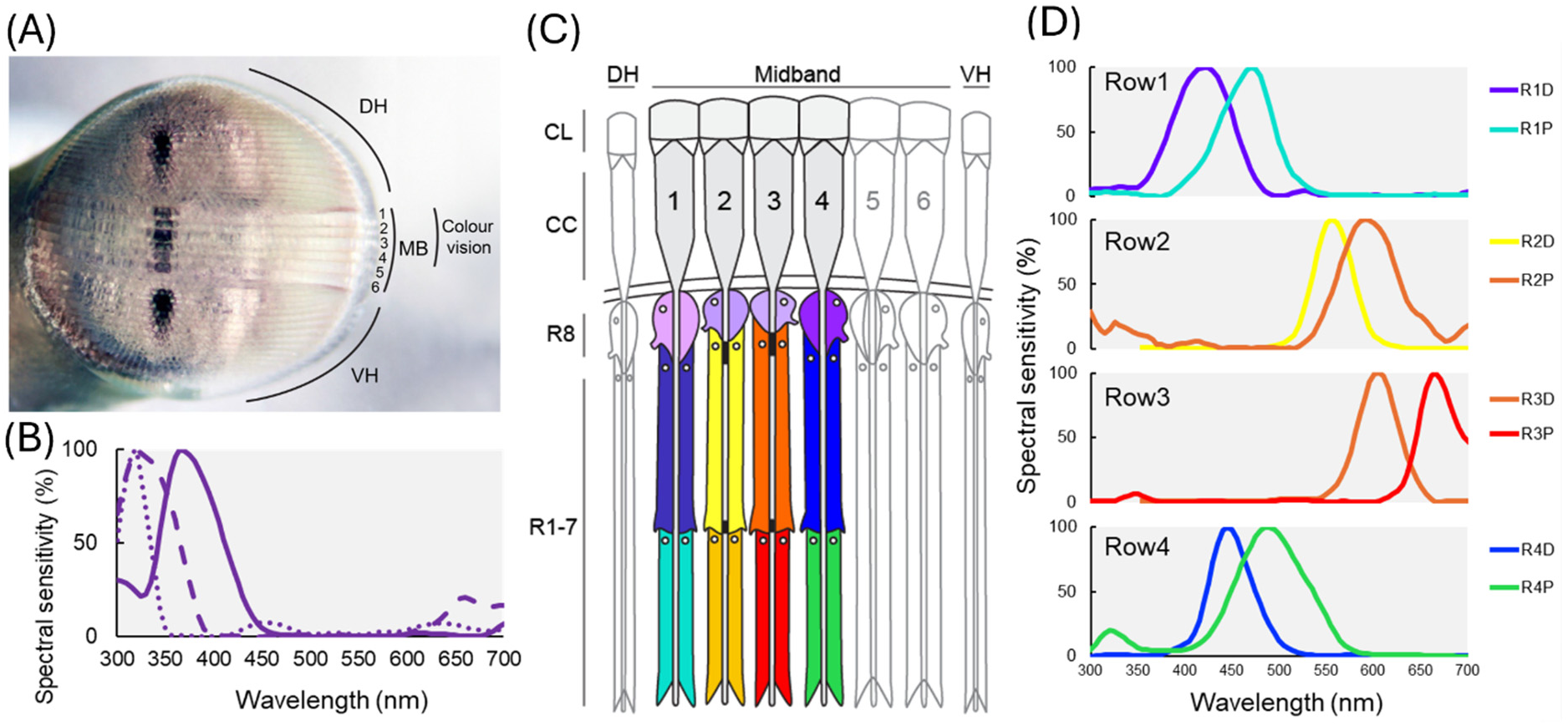
Stomatopod eye structure and spectral sensitivities. (A) Eye of *H. trispinosa,* showing the dorsal/ventral hemispheres (DH/VH) and the six midband (MB) rows. Photo credit: Justin Marshall. (B) Spectral sensitivities of R8, adapted from Thoen et al. (2014) (C) Illustration of the sagittal section of midband rows 1-4, comprising three tiers with 12 spectral sensitivities. CL, corneal lens; CC, crystalline cone; DH, dorsal hemisphere; VH, ventral hemisphere. The illustration was adapted from Marshall et al. (2007). (D) Normalised spectral sensitivities of the two proximal tiers of R-7 in rows 1-4. D: distal; P: proximal. The figures are adapted from Thoen et al. (2014, 2017).

Previous behavioural studies by Marshall et al. (1996) using grey-card-style tests demonstrated that stomatopods possess true colour vision. However, the mechanism stomatopods use to process spectral information in their polychromatic colour vision remained speculative. Two hypotheses regarding the processing mechanism of stomatopods’ unique colour vision have been proposed: (1) multiple dichromatic colour-opponent system and (2) a colour-binning or barcode system (Fig. 2). The multi-dichromatic processing hypothesis suggests that rows 1-4 function as four parallel dichromatic systems, where the spectral information is compared within each row to encode colour information in different spectral ranges; a schema generally similar to other colour vision systems (Fig. 2A) (Marshall, 1988; Cronin and Marshall, 1989b; Marshall and Land, 1993; Marshall et al., 1996; Osorio et al., 1997; Cronin and Marshall, 2001). This hypothesis was initially proposed based on the tiered arrangement of the R1-7 in rows 1-4, which potentially facilitates the comparison of signals perceived from two subsets in a row. This idea is also supported by the ommatidial cell identity and neuronal organisation of their first optic lobe - the lamina ganglionaris (Strausfeld and Nassel, 1981; Kleinlogel et al., 2003; Kleinlogel and Marshall, 2005; Thoen et al., 2017a). Located beneath the retina, the lamina is where the axons of R1-7 terminate, with the terminals of each ommatidium project into individual processing units termed lamina cartridges. The axons of R1-7 terminate at two separate layers in the lamina and form connections with primary processing cells in the lamina, the monopolar cells. Signals carried by these axons are then transmitted to monopolar cells for processing or relaying to the next optic lobe. Such neuronal construction closely resembles the polarisation opponent processing system used for comparing horizontal and vertical polarised signals in many crustaceans as well as the hemispheric regions of the stomatopod eye (Fig. 2B) (Strausfeld and Nassel, 1981; Marshall et al., 1991; Glantz and Miller, 2002; How and Marshall, 2014). Hence, it has been proposed that colour processing in stomatopods may have simply adopted this retinal wiring to compare the different spectral signals perceived by the two subsets of R1-7 in rows 1-4.

**Figure 2.**
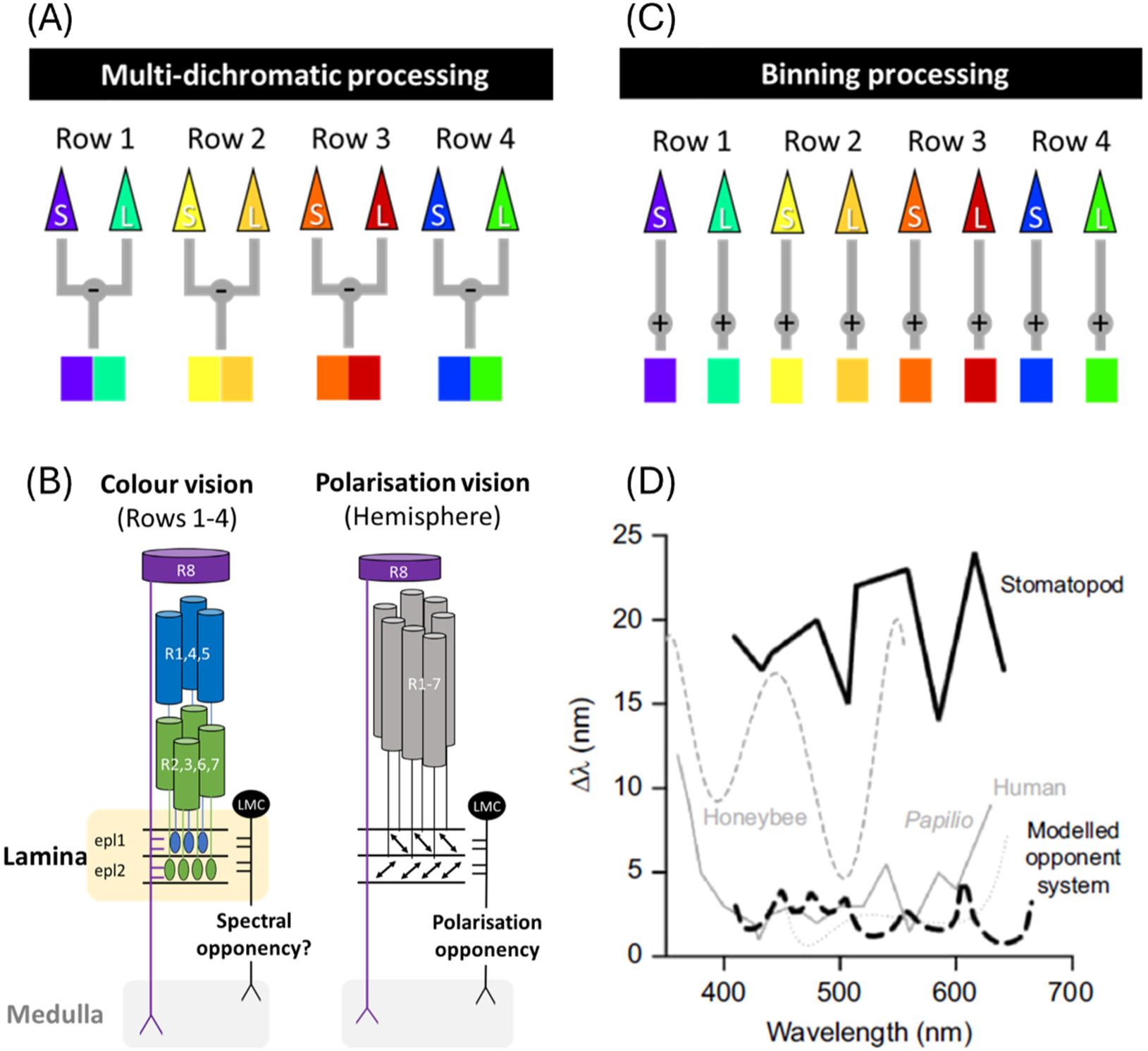
Spectral discrimination curves and schematic diagrams of two processing hypotheses for stomatopods colour vision. (A) Diagram of multi-dichromatic processing, illustrating opponent processing within each spectral-sensitive row. (B) The left diagram illustrates the proposed spectral opponent processing scheme of in midband rows 1-4, using row4 as example. The right diagram shows the polarisation opponent processing scheme in the hemispheres. Epl: external plexiform layer. (C) Binning processing scheme. (D) Spectral discrimination abilities of stomatopods and other animals, showing stomatopods have a coarse spectral resolution compared to the modelled results and other animals. Figure is adapted from Thoen et al., (2014).

Conversely, the binning (barcode) processing system posits that colour perception in stomatopods relies on the activation pattern of the 12 channels, with the channel exhibiting the stronger signal determining the colour (Fig. 2C) (Thoen et al., 2014; Marshall and Arikawa, 2014). This hypothesis was supported by the unexpected low spectral resolution observed in wavelength discrimination tests (Fig. 2D) (Thoen et al., 2014). Having up to 12 spectral classes, stomatopods were expected to possess an exceptionally fine (1-3nm) spectral resolution. However, when animals compared increasingly spectrally close light sources, they were only capable of discriminating stimuli with a 15-25 nm difference (Thoen et al., 2014). Such spectral resolution is worse than that of other visual systems tested in similar ways, despite those systems having fewer spectral receptors (Koshitaka et al., 2008; Thoen et al., 2014). These include visual systems of various animals such as honeybee (5 - 20 nm, von Helversen, 1972), goldfish (3 - 20 nm, Neumeyer, 1986), human (1 - 8 nm, De Valois and Jacobs, 1968) and butterfly (<1 −15 nm, Koshitaka et al., 2008) (see Koshitaka et al. (2008) for original diagram).

This rather surprising finding increased evidence that stomatopods possess a unique colour vision system. Thoen et al. (2014) suggested that stomatopod may bin the spectral information into 12 spectral channels, resembling a diode array system that builds up spectral content from all channels of the array and creates an excitation pattern across their visual spectrum. Such a pattern analyser, also referred to as digital analyser or linear frequency analyser, invokes comparison with the way the cochlea examines auditory space in the mammalian ear and is thought to enable efficient spectral recognition as appose to fine discrimination in stomatopods’ colour vision (Marshall et al., 1997, Thoen et al., 2014; Marshall and Arikawa, 2014).

While the concept of the binning system is intriguing for stomatopod spectral processing and possibly the first of its kind, it does not necessarily exclude the possibility of having other processing methods. Recent study conducted by Streets et al. (2022) tested the colour processing system of stomatopods further across different spectral zones with the experiment built upon one specific result from Marshall et al. (1996). Marshall et al. (1996) showed that the stomatopod *Odontodactylus scyllarus* could distinguish various colour, including green, yellow, red, from grey stimuli, but not blue. This discrepant result in blue was suggested to be linked to the unsaturated blue stimuli used in the tests, resulting in low chromatic contrast compared to the grey stimuli for the stomatopods’ visual system.

To further investigate the opponency and binning hypothesis, Streets et al. (2022) set out to examine if stomatopods indeed are unable to distinguish unsaturated colours with grey stimuli. They hypothesised that if their visual system relies solely on a binning system for spectral processing, low-saturated colours over the whole spectrum (not just in the blue) versus grey would appear indistinguishable, as they generate similar activation patterns across the spectral channels. Interestingly, their experiments showed that stomatopods are capable of discerning between low-saturated colours and grey stimuli, including in the blue. This implies stomatopods colour vision may not simply be established on a binning processing, instead, some forms of opponent comparison must be at play to allow them to differentiate between low-saturated colours and greys.

Additionally, the results from Streets et al. (2022) also shed new light on the findings in Marshall et al. (1996). The discrepancy of the two studies is thought to be attribute to several possible factors, including: a) The use of different species in these two studies, b) A post 1996 discovery of how rapidly the stomatopod colour sense (and indeed other animal colour vision) may degrade in artificial room lighting, a factor highlighted by subsequent studies in mantis shrimp (Cronin et al., 2001; Cronin et al., 2002; Cronin and Caldwell, 2002; Cheroske et al., 2003; Cheroske and Cronin, 2005) and also in reef fish (Carleton et al., 2020). As a result, the lack of discrimination ability in Marshall et al. (1996) is suggested to have been influenced by the time animals spent in captivity in a short wavelength-poor room light, a factor unknown at that time. Despite these inconsistencies, it is interesting to note that the results from Marshall et al. (1996), Streets et al. (2022) and indeed those described below indicate that blue may hold a special function or significance for stomatopods.

While previous studies have provided valuable insights into the spectral processing mechanism of stomatopods, there has been a lack of direct behavioural evidence regarding the hypothetical multiple dichromatic opponent processing in their colour vision. Therefore, the purpose of this study was to explore this further using a behavioural approach. We applied modified von Frisch grey card experiments (Frisch, 1914; Kelber et al., 2003; Streets et al., 2022) to test their colour vision under natural light and three different coloured illuminations, blue, green and red, with restricted spectra. Under coloured illumination, the spectra of the stimuli become a challenge for the binning system as they would produce comparable excitation patterns. This would make it difficult to distinguish between colour and grey stimuli. However, with opponent processing, different spectral classes could yield distinct outputs, enabling discrimination even under coloured illumination. We, therefore, hypothesised that if stomatopods possess an opponent processing system in their colour vision, they would be capable of identifying colours from greys under coloured illuminations.

## Materials and Methods

### (1) Animal collection

Stomatopods, *Haptosquilla trispinosa*, were collected from shallow reef areas on Lizard Island Research Station (LIRS) at a depth range of 0 to 5 m (GBRMPA permit no. G17/38160.1). Following collection, each individual was placed in behavioural chambers for a period of 1-2 day for acclimation. Each chamber contained a grey PVC tube designed as a burrow with front and back lids. The front lid had a drilled hole serving as the burrow’s entrance. To mimic their natural habitats, fresh sand sourced from their original environment was added to the chamber, filling it to approximately one-third of its depth to create a sandy base. To avoid degradation of their colour vision, the experiments were performed only on freshly caught stomatopods.

This study comprised two separate experiments; the first was conducted from July 19^th^ to August 12^th^, 2021, and the second from February 19^th^ to March 5^th^, 2022. Both experiments took place at the outdoor bench area at LIRS, sheltered beneath clear polycarbonate roofs. While some UV is blocked from above by this roofing, UV still reaches the experimental arena from the sides. As the experiments described here are not in UV wavelengths, this slight reduction was assumed irrelevant.

### (2) Stimulus and illumination design

A modified von Frisch grey card experiment (Fig. 3A) (Frisch, 1914; Kelber et al., 2003; Templin, 2017; Streets et al., 2022) was employed to investigate stomatopods’ ability to distinguish colour from greys under natural light and various illumination conditions. White plastic cable ties (Crescent, 2.5×2.5×10 mm) were used to present different stimuli to stomatopods. The size of the cable ties was specifically chosen for *Haptosquilla trispinosa* to allow easy grasping and transportation back to their burrows. Each cable tie was fitted with colour or ND filters on the front surface using white double-sided tape to create colour (target) and grey (distractors) stimuli. The reflectance of the stimuli was shown in Figure. 3C. The colour filters used included blue (≈ 360-550 nm; LEE Filters #183), green (≈ 500-560 nm; LEE Filters #139) and red (>610 nm; LEE Filters #027). Four intensities of grey distractors were created using ND 0.15 (LEE Filters #209), 0.3 ND (LEE Filters #209), 0.6 ND (LEE Filters #210) and 0.9 ND (LEE Filters #211).

**Figure 3.**
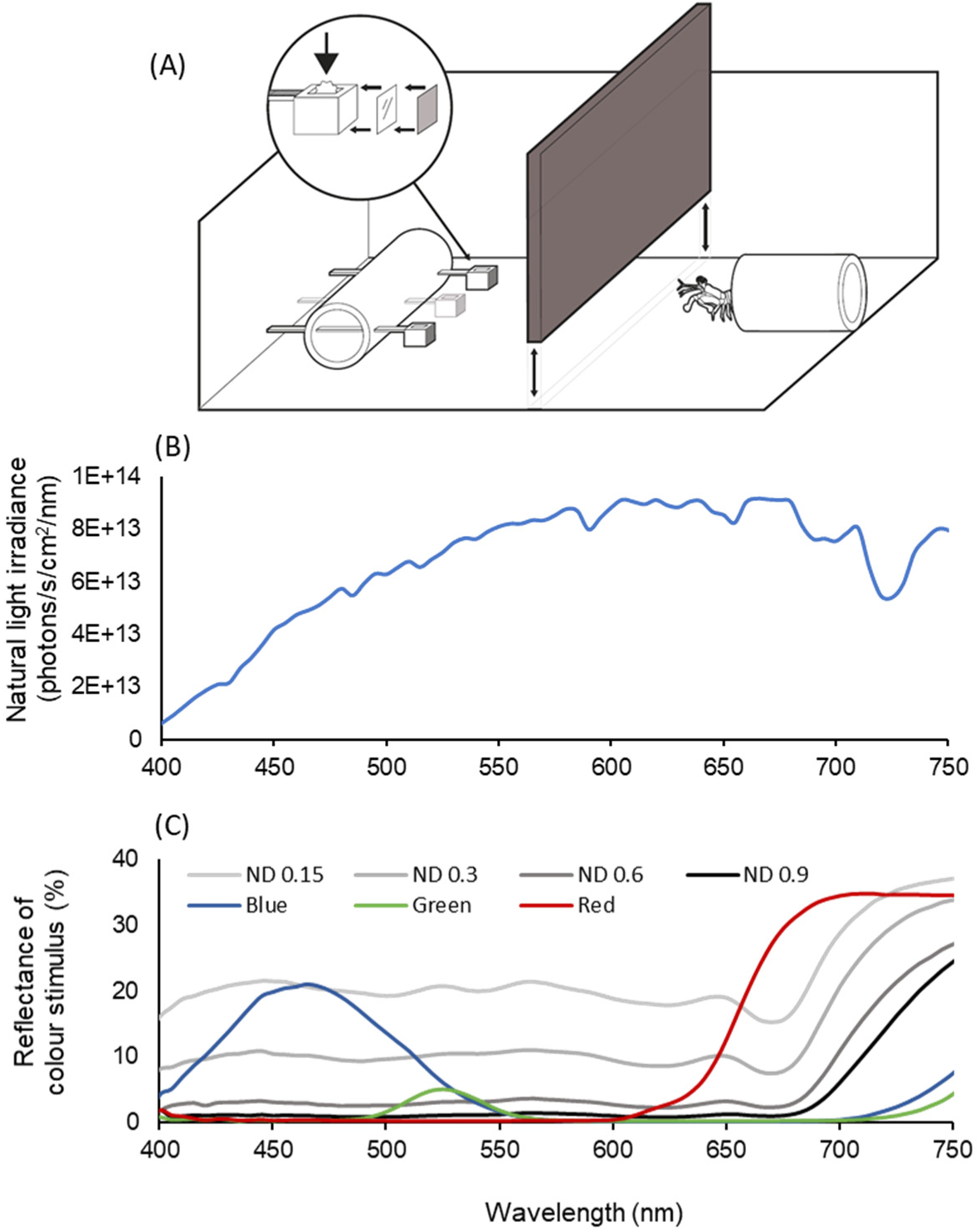
Experiment design of the grey card experiment. (A) Diagram adapted from Streets et al. (2022), showing the behavioural chamber for colour discrimination tests. During training, food reward was placed inside the camber of the target cable ties. (B) Irradiance spectrum of the natural light illumination at Lizard Island Research Station, measured at where the experiments were conducted. (C) Reflectance spectra of colour and grey stimuli, measured from the surface of cable ties.

During both training and testing trials, three stimuli, consisting of one target and two distractors, were presented to the stomatopods. An illustration of the experiment set up is shown in Figure 4. Food rewards, a small piece of shrimp meat, were exclusively placed in the ratchet end cavity of the colour cable ties (targets), while the grey cable ties (distractors) remained empty and uncontaminated. In addition, customized cable tie holders, crafted from white PVC tubes perforated with three holes aligning horizontally at each side, were utilised to display the cable ties with consistent distance and height throughout the experiments. The displaying distance between stimuli and the animals ranged from 5 to 8 cm, slightly longer than the animals’ body length, compelling them to fully exit their burrows when making a choice.

**Figure 4.**
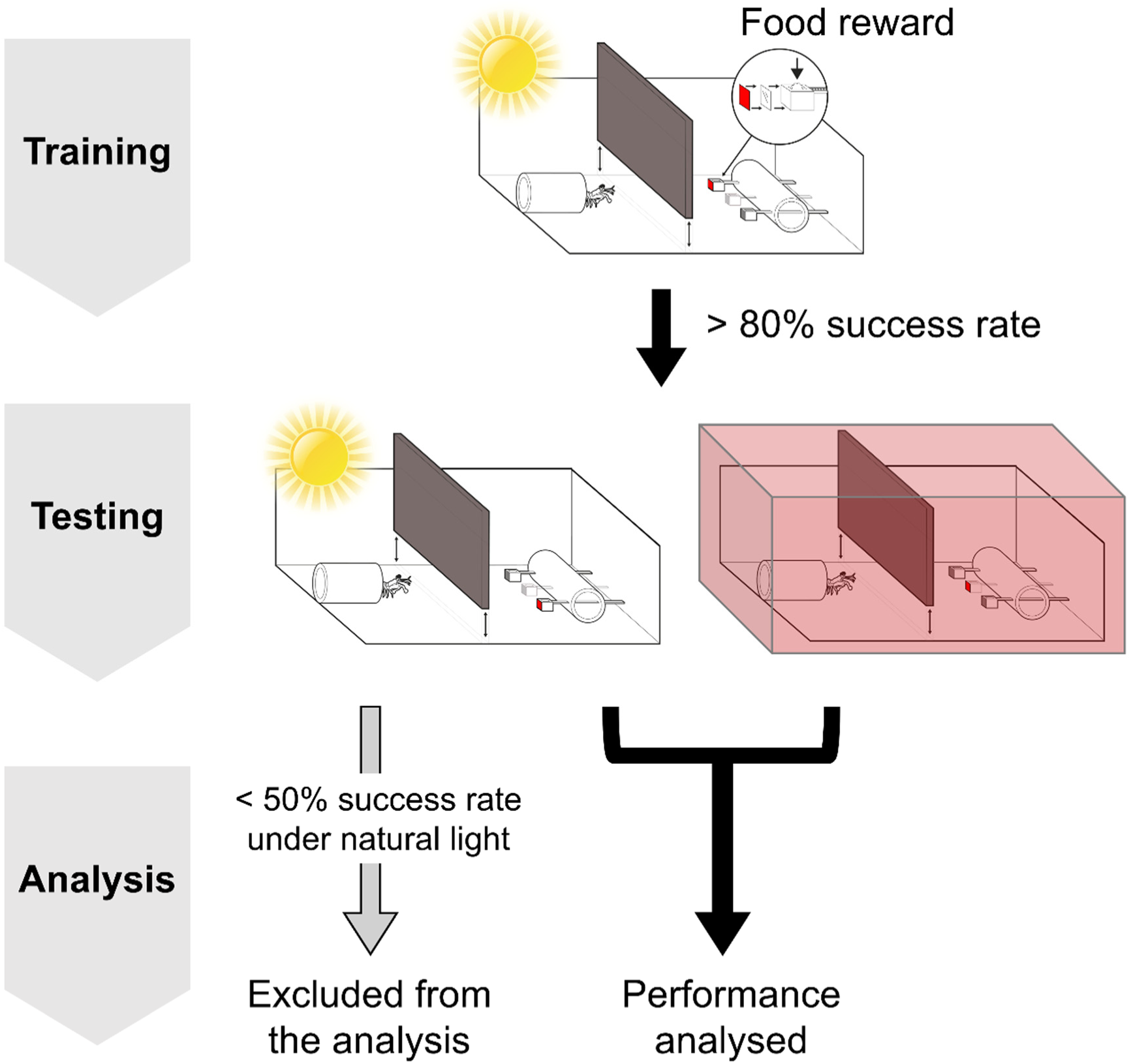
Procedure of the colour discrimination ability experiment. During training, food reward was placed in the colour cable tie while the grey cable ties remained empty. Animals that passed 80% success rate for 5 consecutive trials were moved to the testing stage. At the testing stage, stomatopods’ colour discrimination abilities were tested under natural light and coloured illumination. Individuals with success rate lower than 50% under natural light were excluded from the experiment as the low success rate under natural light indicated that they failed to associate the reward with visual stimuli. For those individuals that were able to maintain high performance under natural light, their performances under coloured illumination were analysed to examine their colour discrimination abilities under coloured illuminations.

In terms of the light condition, the first experiment was conducted under natural light and coloured illuminations, blue, green, and red lights. To alter the lighting within each behavioural chamber, customised filter tents were crafted from plastic sheets and applied during coloured illumination tests. The filters used to provide coloured illuminations included blue (≈ 360-590 nm; LEE Filters #172), green (≈ 450-600 nm; LEE Filters #124) and red (>590 nm; LEE Filters #182). The second experiment aimed to testes their colour vision under dimmer natural light and coloured illuminations. For creating dimmer coloured lighting, double-layered colour filters were used to reduce the light intensity, whereas natural light intensity was adjusted using ND filters, including 0.3 ND (LEE Filters #209), 0.6 ND (LEE Filters #210) and 0.9 ND (LEE Filters #211). The transmittance of these filters is illustrated in Figure. 5A and Figure. 6A.

**Figure 5.**
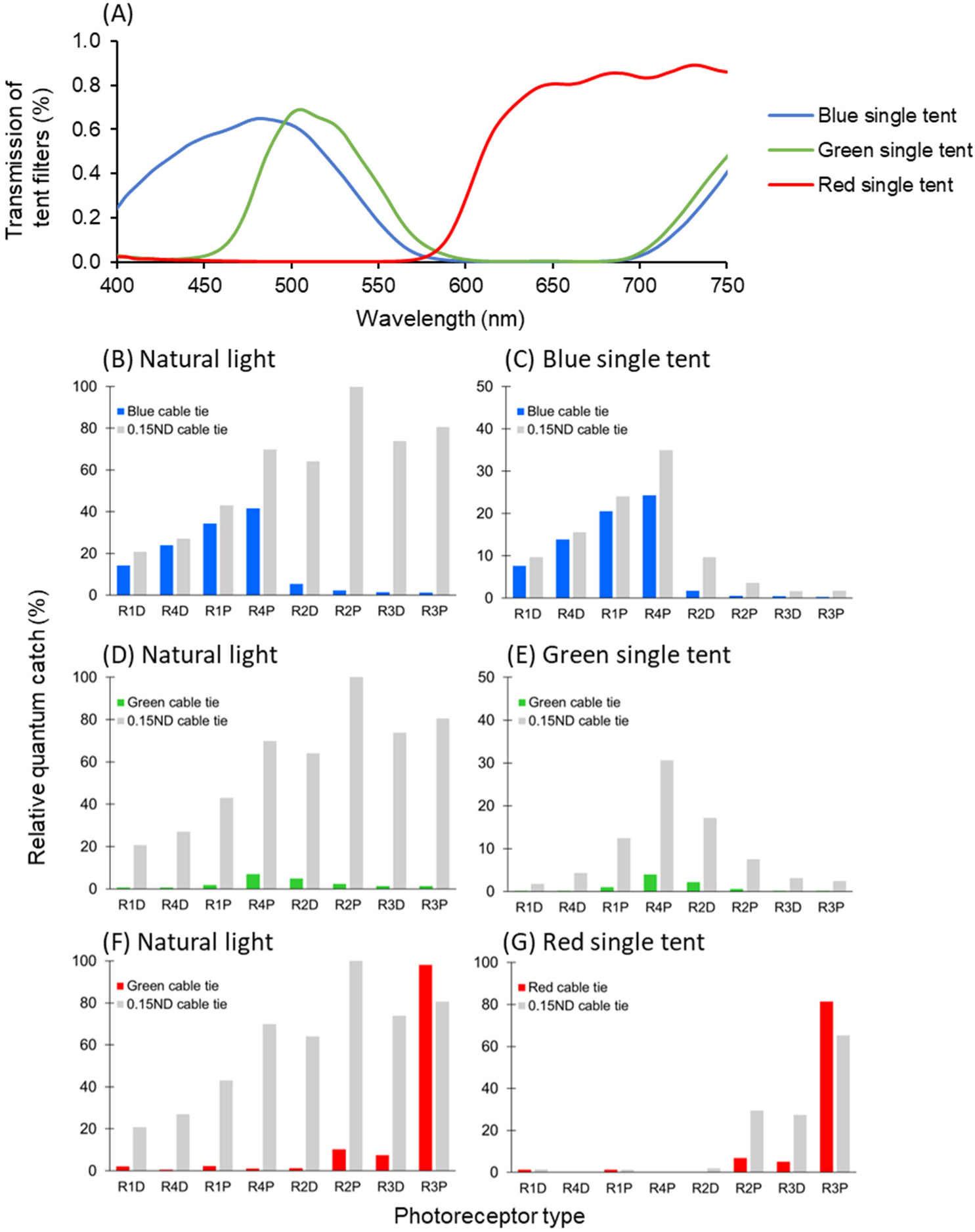
Transmittance of the tent filters of Experiment 1 (single-layered tent) and the relative photon catch for each photoreceptor under natural light and coloured illuminations. (A) Transmittance of the filters used to change the illumination. (B) (C) Normalised quantum catch of the receptors when viewing blue and grey (ND 0.15) stimuli under natural light and blue illumination. (D) (E) Normalised quantum catch of the receptors when viewing green and grey (ND 0.15) stimuli under natural light and green illumination. (F) (G) Normalised quantum catch of the receptors when viewing red and grey (ND 0.15) stimuli under natural light and red illumination. The order of the photoreceptor type is arranged according to the spectral sensitivities of receptors, numbers indicate the row number of rows 1-4; D: distal; P: proximal. All photons catch values were normalised to the highest photon catch value observed from R2P when stimulated by grey (ND 0.15) under natural light.

**Figure 6.**
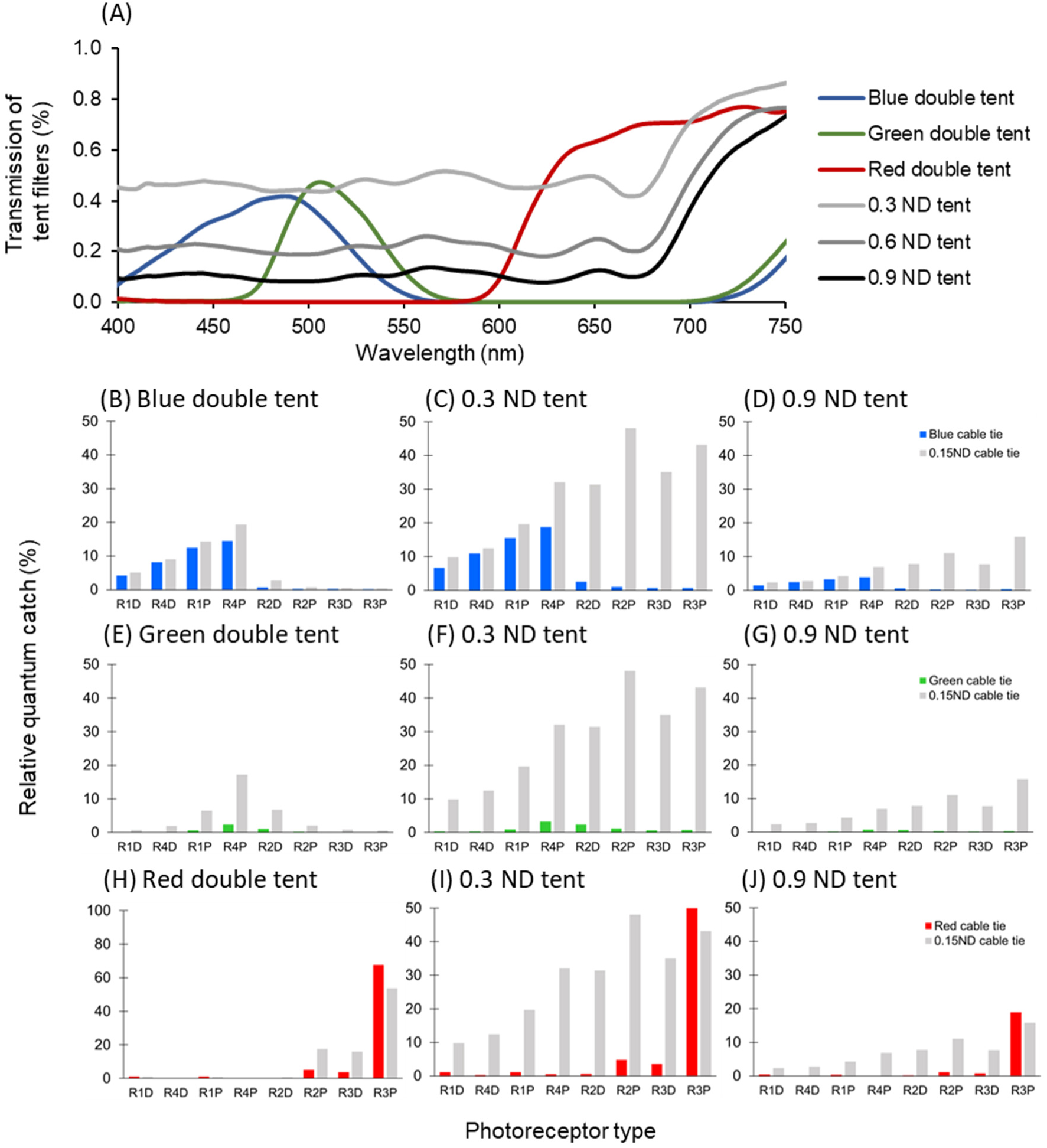
Transmittance of the tent filters of Experiment 2 (double-layered and grey tents) and the relative photon catch for each photoreceptor under different illuminations. (A) Transmittance of the filters used to change the illumination. (B)(C) (D) Normalised quantum catch of the receptors when viewing blue and grey (ND 0.15) stimuli under double-layered blue tent and different ND filter tents. (E)(F)(G) Normalised quantum catch of the receptors when viewing green and grey (ND 0.15) stimuli under double-layered green tent and different ND filter tents. (F)(G) Normalised quantum catch of the receptors when viewing double-layered red tent and different ND filter tents. Note that there were 3 ND filter tents used to modify the natural light intensity, here only showing the quantum catch of 0.3 and 0.9 ND as examples. All photon catch values were normalized to the highest photon catch value observed from R2P when stimulated by grey (ND 0.15) under natural light.

### (3) Experiment design

An established choice test procedure outlined by Templin (2017) and Streets et al. (2022) was followed to train and test stomatopods. The experiment comprised of three stages: priming, training and testing. Each animal was assigned to one of the three colour targets, blue, green or red, for the experiment.

#### Priming

During the priming stage, animals were introduced to a single cable tie of their assigned colour. The cable tie contained food in the ratchet cavity and was placed at one of the three positions (left, middle and right) in the holder. The position of the cable tie varied for different trials with a random order. In each priming trial, animals were given 5 minutes to grasp the cable tie before the trial was aborted. After the trial, cable ties were retrieved, cleaned with fresh water and recycled for future trials. All animals underwent priming trials three times a day (morning, midday and afternoon) for 2-3 days. Individuals that reached an 80% participation rate in the last 5 consecutive trials were allowed to proceed to the training stage. Animals with participation rates lower than 50% were considered unmotivated, potentially due to stress, sickness or moulting. Therefore, these animals were deemed untrainable and replaced by new animals.

#### Training

During the training stage, animals were presented with three stimuli in each trial including one colour stimulus (target) and two grey stimuli (distractors) (Fig 4, Training). Only the colour stimulus contained a food reward in the cable tie, while the distractors remained empty and odourless. The grey stimuli were selected from four intensities of grey stimuli with 0.15, 0.3, 0.6 and 0.9 ND filters. The position and combination of the stimuli were arranged using a pseudo-randomised method, with adjustments made to avoid repetitive colour or grey stimuli occurring at the same position for more than two trials. The stimulus combinations are listed in Table S1 in the Supplementary Appendix.

Animals were given five minutes for each training trial. A correct choice was recorded if their first selection was the colour cable tie. Conversely, it was considered wrong choice if they chose a grey cable tie. Regardless of their choice, animals were permitted to explore any other cable ties within the five-minute time frame.

All training trials were conducted three times a day (morning, midday and afternoon) under natural light. The performance of each individual was assessed after three days of training (9 trials). Individuals with over 80% success rate (correct choice) in the last six consecutive trials were allowed to proceed to the testing stage. Individuals with success rate between 30% to 80% were kept in the training group for more training trials. Those with success rates below 30% were considered untrainable and, therefore, removed from the experiment.

#### Testing

In the testing phase, all colour target and ND distractor stimulus were constructed with new cable ties, carefully kept separate from the training cable ties to avoid any food and therefore olfactory contamination. Similar to the training trials, one colour target and two grey distractors were presented in each testing trial. The combination of the stimulus was also randomised with adjustments to avoid recurrence of the same combination set or target positions. Three trials were conducted each day in the morning, midday and afternoon, respectively. Each test was conducted under either natural light or coloured illumination conditions (Fig 4, Testing; Movie 1). To alter the lighting during the test, a colour tent was applied to fully cover the behavioural chamber throughout the test. Before each trial began, the stimuli were placed into the chamber by slightly lifting the tent. In the meantime, the stimuli were carefully covered by the experimenter’s fingers when being transported into the chamber.

The testing phase commenced with natural light tests on the first day, and coloured illumination tests were introduced from the second day onwards. This gradual transition was to allow individuals to accustomed to the testing procedure. Starting from the second day of testing, one natural light and two colour illumination tests were performed in random order each day for every individual.

In each testing trial, the individuals had up to five minutes to make a choice in each trial. However, the trials ended once a choice was made. A correct choice was marked when the individual grabbed or struck the colour cable tie. After a correct choice was made, a food reward would be delivered to the individual with forceps. On the contrary, if the individual made an incorrect choice, the cable ties would be retrieved, and no food would be given to the animal. Similarly, if the individual did not make any choice during the time frame, the cable ties would be removed, and the individual would not receive food reward.

Additionally, reinforcement trials were incorporated after every six testing trials. Following the same procedure as training trials, these reinforcement trials provided food rewards with cable ties, allowing individuals to explore all options within the designated time frame. Notably, choices made during reinforcement trials were omitted from subsequent data analysis.

### (4) Statistical analysis

The purpose of the analysis was to compare stomatopods’ colour discrimination abilities under both natural light and coloured illumination and determine whether their performances under the two conditions differ.

After the individuals completed each testing trial, the overall performance of each individual was carefully examined before analysis. The colour illumination tests aimed to assess colour discrimination abilities under a restricted spectrum (blue, green, and red), while natural light tests were conducted simultaneously to evaluate the reliability of each individual’s performance. With a 33% probability of randomly choosing the colour stimulus, a threshold of a 50% success rate was established to determine whether an individual was well-trained and capable of maintaining their performance throughout the testing phase (Fig 4, Analysis). To ensure the reliability of the individuals’ performance, only animals demonstrating a success rate higher than 50% under natural light conditions were included in the data analysis. Individuals with poor performance (below 50%) under natural light conditions were considered to have failed in training, possibly due to factors such as forgetting tasks or experiencing physical issues like stress, sickness, or moulting. In such cases, their performance under coloured illumination was regarded as unreliable and subsequently excluded from the analysis.

All analyses were performed in R studio (v4.1.2). Chi-square independence tests were employed to evaluate colour discrimination ability, comparing observed and expected counts of choices in three-choice tests with one target and two distractors. Expected counts for correct and incorrect choices were calculated as [total counts] × ⅓ and [total counts] × ⅔, respectively.

Additionally, Generalized Mixed Models (GLMM) were executed using the lme4 package to explore the relationship between various factors (individuals, light treatments, or target positions) and choice outcomes. The model, applying a binomial distribution with a ‘logit’ link function, considered animal choices during tests as the variable of interest. Individual ID and sex were treated as random effects due to multiple tests per individual, while light treatment, distractor combination, and position of choice were considered fixed effects. However, sex, not significantly contributing to explanatory power, was excluded from the final models.

### (5) Quantum catch of different photoreceptor types

When examining different stimuli under various illuminations, the activation levels of photoreceptors were quantified using the formula:

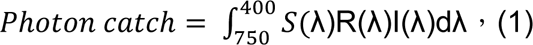

where S(λ) represents the spectral sensitivity of the receptors of *H. trispinosa*, determined by the intracellular recording results (Thoen et al., 2017b); R(λ) is the reflectance of the stimuli; I(λ) is the photon number distribution of the illumination, and λ denotes the wavelength. Figure. 5B-G illustrate the photon catches of receptors activated under different conditions in Experiment 1, while Figure. 6B-J show the photon catches of receptors in Experiment 2.

## Results

### (1) Experiment 1 - Colour Discrimination Ability under Colour Illumination

In Experiment 1, a total of 69 stomatopods were trained, including 27 in the blue group, 20 in the green group, and 22 in the red group. Among these, 20 exhibited reliable performance under natural light conditions throughout testing, comprising 4 in the blue group, 8 in the green group, and 8 in the red group. Figure. 7 illustrates the success rates under natural light and coloured illuminations. For the blue group, individuals displayed proficiency in identifying blue stimuli from greys under natural light with a success rate of 76% (N_individuals_ = 4; N_trials_ = 25). However, their performance declined to 40% when tested under blue illumination (N_individuals_ = 4; N_trials_ = 20). Statistically, it was not significantly different from the random choice rate of 33% (Chi-square test: p = 0.3291). Moreover, GLMM analysis indicated a significant decrease in their correct choice rate under blue tents compared to natural light conditions (GLMM, p < 0.05), suggesting that blue-trained animals struggled to distinguish blue stimuli from distractors under blue illumination.

**Figure 7.**
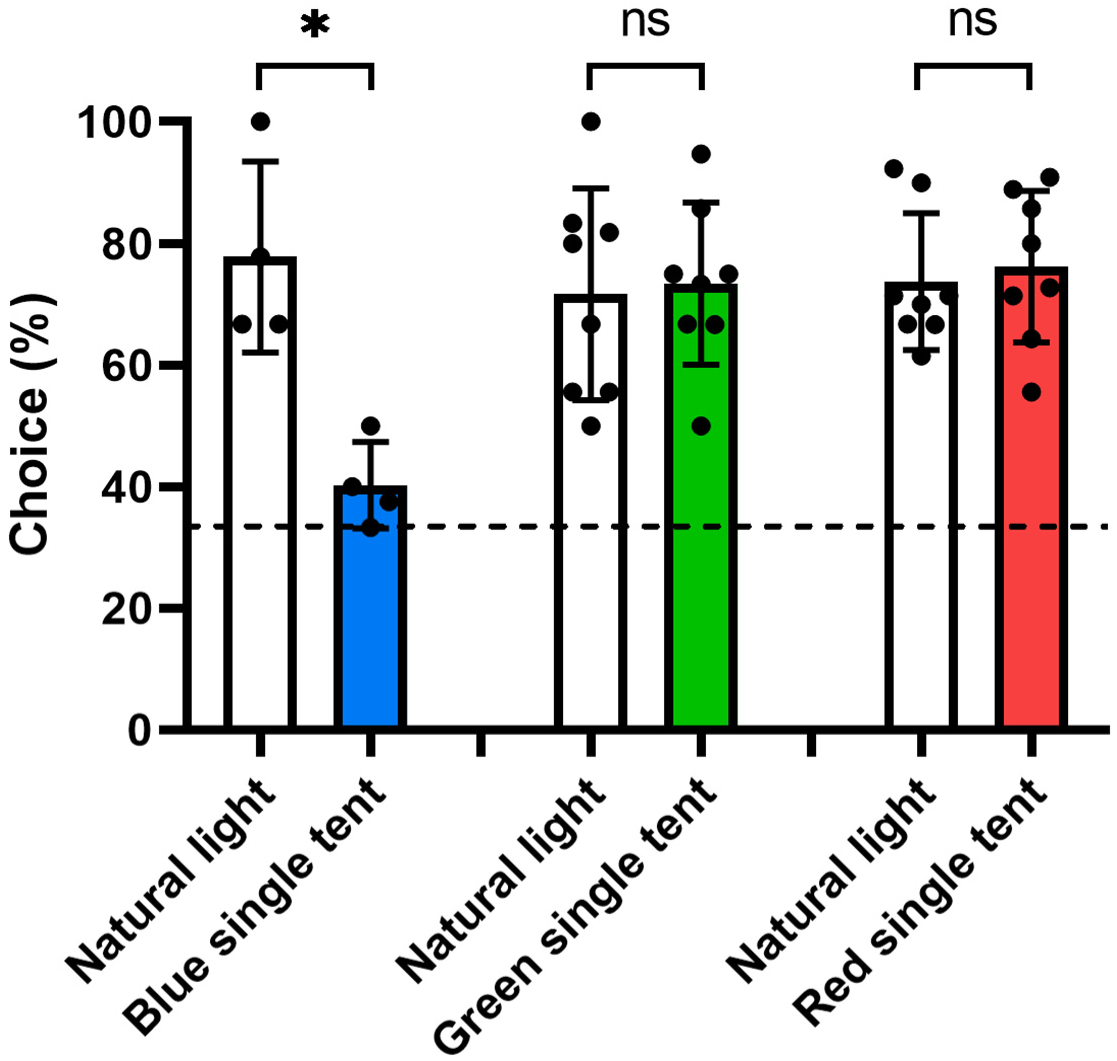
Colour discrimination abilities of *H. trispinosa* under natural light and different coloured illuminations. Histogram shows the success rates of blue, green and red trained *H. trispinosa* when tested under natural light and coloured illumination. Each dot indicates the average performance of an individual. Dashed line represents the random choice probability (33.3%) from the three-choice test. Significance: *: P < 0.05, ns: no significance.

In contrast, individuals in the green and red groups showcased the ability to distinguish colours from greys even under coloured illuminations. The green group, with a success rate of 71.08% under natural light (N_individuals_ = 8; N_trials_ = 83), maintained a high success rate of 75.7% under green illumination (N_individuals_ = 8; N_trials_ = 107). Similarly, the red group, having a 74.73% success rate under natural light (N_individuals_ = 8; N_trials_ = 91), sustained a success rate of 73.74% under red illuminations (N_individuals_ = 8; N_trials_ = 99).

These choice rates significantly deviated from the probability of random choices (Chi-square test: green, p < 0.0001; red, p < 0.0001) and did not significantly differ from the performance tested under natural light (GLMM, pGreen = 0.5205; pRed = 0.956). These findings indicate a consistent ability to discern colours from greys, irrespective of changes in illumination, emphasizing their capability to conduct colour vision with only 2-3 spectral receptors operating under restricted spectra.

### (2) Experiment 2 - Colour Discrimination Ability under Dimmer Illuminations

In this experiment, 40 individuals were primed and trained. Only 15 showed reliable performance under natural light tests and were included in the data analysis, consisting of 4 individuals from the blue group, 6 from the red group, and 5 from the green group.

For the blue and red groups, stomatopods’ colour discrimination ability declined under dim coloured illumination compared to their performance under natural light (Fig. 8). The blue group, which could discriminate blue from greys with a 71.86% success rate under natural light (N_individuals_ = 4; N_trials_ = 32), experienced a significant drop to 29.41% under the double-layer blue tent (N_individuals_ = 4; N_trials_ = 34). This success rate was significantly lower than that under natural light (GLMM, p < 0.0001) and not significantly different from random choice (33.3%) (Chi-square test: p = 0.9403), indicating their inability to discriminate blue targets from greys under dimmer blue illumination.

**Figure 8.**
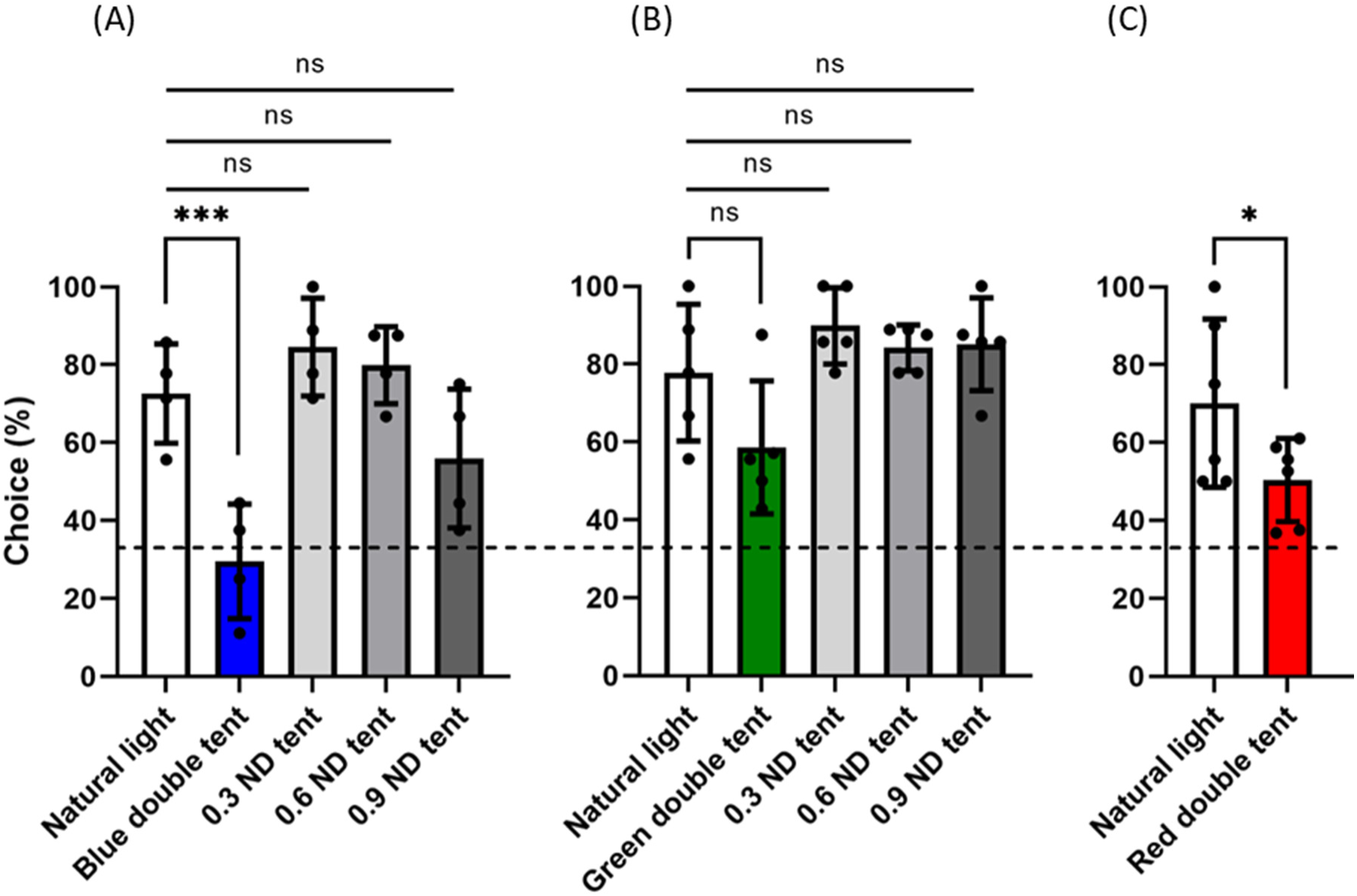
Colour discrimination abilities of *H. trispinosa* under natural light and different dim illuminations. (A) Performance comparison under double-layered blue tent and varying natural light intensities. (B) Performance comparison under double-layered green tent and varying natural light intensities. (C) Performance comparison under double-layered red tent and varying natural light intensities. Each dot indicates the average performance of an individual. Dashed line represents the random choice probability (33.3%) from the three-choice test. Significance: *: P < 0.05, **: P < 0.01; ***: P < 0.001; ns: no significance.

Similarly, the red group exhibited a decline in performance under dimmer red light. While having a success rate of 68.97% under natural light (N_individuals_ = 6; N_trials_ = 58), their success rate reduced to 50.70% under double-layer red tent (N_individuals_ = 6; N_trials_ = 71). Although this success rate was significantly different from random choice probabilities (33.3%) (Chi-square test: p < 0.001), it was significantly lower than their performance under natural light condition (GLMM, p < 0.05).

However, the green group demonstrated a high success rate under dim-coloured illumination. There was no significant difference between the success rate under natural light (78.57%, N_individuals_ = 5; N_trials_ = 42) and double-layer green tent (58.97%, N_individuals_ = 5; N_trials_ = 39) (GLMM, p = 0.6).

In the ND tent tests, blue and green groups showed exhibit similar, achieving high success rates under varying natural light intensities. For the blue group, the success rates under 0.3 ND, 0.6 ND and 0.9 ND tents were 84.38% (N_trials_ = 32), 79.41% (N_trials_ = 34), and 55.17% (N_trials_ = 29), respectively. No significant differences were found between the success rates under natural light and any ND tent (GLMM, pND0.3 = 0.22, pND0.6 = 0.46, pND0.9 = 0.20).

Likewise, the green group maintained a high success rate of over 80%, even with the dimmest light treatment. The success rates under different ND tents were 89.19% for the 0.3 ND tent (N_trials_ = 37), 84.09% for the 0.6 ND tent (N_trials_ = 44) and 86.84% for the 0.9 ND tent (N_trials_ = 38). No significant differences in their success rates between natural light and any ND tent treatments in the green group (GLMM, pND0.3 = 0.21, pND0.6 = 0.51, pND0.9 = 0.34).

It is important to note that no ND tent treatments were applied to the red group due to the ND filters’ non-neutral filtering property in the long wavelength range. Although these filters exhibit effective neutral filtering effects in the short- and medium-wavelength ranges, they are unable to attenuate light effectively in the red spectral range (over 680 nm), making them unsuitable for the purpose of the test (Fig. 6A). Therefore, the ND tent test for the red group was aborted.

## Discussion

### (1) Behavioural evidence of spectral processing in stomatopods’ colour vision

Numerous studies have investigated the spectral processing system underlying stomatopods’ intricate colour vision (Marshall, 1988; Cronin and Marshall, 1989b; Marshall et al., 1991; Marshall et al., 1996; Osorio et al., 1997; Cronin and Marshall, 2001; Thoen et al., 2014; Streets et al., 2022). This study specifically explores the hypothesis of multi-dichromatic opponent processing in stomatopods’ colour vision by conducting modified colour discrimination tests on *H. trispinosa*.

Results revealed that stomatopod *H. trispinosa* can discriminate colour from grey stimuli in the green and red spectral windows, though not under blue. Despite the discrepancy observed under blue illumination, the high success rates under green and red illuminations provide compelling evidence that the stomatopods can achieve colour vision with visual inputs from only a few rows in the midband rows 1-4. This finding reinforces the likelihood of opponent processing in stomatopods’ colour vision system, as distinguishing coloured and grey stimuli under coloured illumination would be implausible with the sole use of a binning system. This result supports the hypothesis proposed by Streets et al. (2022) that opponent and binning processing systems may coexistence in stomatopods’ colour vision.

However, how opponent processing operates in stomatopods’ 12-channel colour vision? Can each row function as an independent dichromatic colour vision system? While different light treatments we applied allow activation of different sets of photoreceptors, the ideal scenario is to limit the function to only one row to examine opponent processing in that specific row. However, none of the light treatments used or indeed known manage to restrict activation to a single row, owing to the intertwined spectral sensitivities of rows 1 and 4, and the overlapping of the spectral sensitivities of different rows (Thoen et al., 2017b). Blue (400-600 nm) and green (450-600 nm) illuminations activated rows 1, 2 and 4 to varying extents. Red illumination (550-750 nm) activated the least classes of photoreceptors, with the spectral range permitting full activation of only row 3 and partial stimulation of row 2 receptors.

Nevertheless, the experiments yielded valuable insights into the opponent processing of stomatopods’ colour vision. Firstly, achieving colour vision with only 2-3 rows implies the processing between different rows may be somewhat independent. This idea finds supports in the neural architecture and connectivity of rows 1-4. These four rows exhibit segregated construction at the retina level, forming four distinct parallel channels. Furthermore, the receptor axons of each row project parallelly into the lamina and terminate at individual processing units of each row in the lamina. These findings substantiate that rows 1-4 function as four independent processing channels, with information perceived from each row being kept separate. Secondly, stomatopods demonstrated colour vison under red illumination, with only row 3 operating in full capacity while row 2 partially activated. As it is suggested that there is minimal comparison or processing between rows, it is inferred that row 3 provided the critical information for distinguishing colour under the red illumination. These findings suggests that stomatopods could most likely perceive colour vision primarily within a single row, through the comparison of signals perceived from the two subsets of R1-7. Overall, given the identical construction scheme across all rows in rows 1-4, it is presumed that the same dichromatic opponent processing is applied in each row.

On the other hand, Experiment 2 investigated the colour discrimination ability of stomatopod *H. trispinosa* under dim natural light and coloured illumination. The dim coloured illumination treatment was constructed by applying two layers of colour filters, while the dim natural light treatments included different filters, 0.3 ND, 0.6 ND, and 0.9 ND. The dimmest natural light condition (0.9 ND) resulted in the lowest photon catch for their visual system, lower than double-layered colour tents. Notably, despite having a higher overall photon catch level, colour discrimination ability under dim coloured illuminations tended to decline more than that under dim natural light conditions. This implies that while rows 1-4 may function as multi-dichromatic system and can obtain colour vision within a single row, the full capacity of rows 1-4 may still be more advantageous for colour discrimination, particularly in dim light conditions. However, whether this advantage is attributed to the activation of all spectral rows, or the incorporation of luminance information remains unknown. This experiment was preliminary, and future studies with careful brightness control are needed to understand the role of light intensity in colour discrimination under various spectral conditions.

Overall, our findings provided the first direct behavioural evidence of opponent processing in colour vision. This discovery strongly supported the multiple colour-opponent system hypothesis (Marshall, 1988; Cronin and Marshall, 2001; Marshall et al., 1996; Osorio et al., 1997), suggesting that each row in the midband rows 1-4 can operate as an independent processing channel. However, it is important to highlight that, while our findings provide compelling behavioural evidence for spectral opponency in stomatopods, they do not exclude the existence of another form of colour vision, such as the binning system (Streets et al., 2022). This perspective helps in clarifying the lack of fine spectral discrimination in stomatopods observed in Thoen et al. (2014). Their coarse colour resolution supports the idea of a combination of binning and opponent processing mechanism in the stomatopod colour vision, a point discussed in greater detail in subsequent sections.

### (2) Deciphering the discrepancy: Behavioural variances with blue stimuli

Our experiment examined the colour discrimination ability of stomatopod *H.trispinosa* under three spectral zones: blue, green and red. While the green and red group exhibited significant success in discerning colour from grey stimuli under coloured illuminations, the blue group, in contrast, failed to make this distinction under blue illumination. It seems unlikely that stomatopods could not perceive colour under blue illumination as they do under green and red illuminations. The blue illumination used in the tests covered the widest spectrum range and activated the greatest number of photoreceptor classes. Hence, in theory, blue group is supposed to perform similarly to the other groups.

Interestingly, the blue group not only failed in colour discrimination under blue illumination tests, but also exhibited distinct behaviour compared to the other two groups. To begin with, it required a larger number of individuals to undergo training to reach the threshold of over 8 individuals needed for testing. Additionally, while all groups demonstrated their abilities under natural light during testing, the blue group struggled to maintain their performance. The correct choice rate of blue group was 53.6% when tested under natural light (N_individuals_ = 9, N_trials_ = 69), which is much lower than the green (72.72%, N_individuals_ = 9, N_trials_ = 88) and red groups (74.73%, N_individuals_ = 8, N_trials_ = 91). Subsequently, five individuals in the blue group were excluded from the analysis due to poor performance (correct choice rate under 50% under natural light condition), in contrast to one in the green group and none in the red group. This resulted in a much lower trainability for the blue group (i.e. the ratio of the individuals trained and those meeting the threshold for analysis) compared to the green (40%) and red groups (36.36%). Lastly, the blue group showed about three times higher rates of no-choice instances during testing both natural light and coloured illuminations compared to the green and red groups. For more details of the performances of each group, see Table. S2 in the Supplementary Appendix. These observations collectively indicate that blue was a more difficult cue to learn compared to the other two colours.

As mentioned in the introduction, this is not the first time blue has posed an interesting challenge to stomatopods. Marshall et al. (1996) previously observed difficulties in stomatopod *O.scyllarus* in discerning blue from greys, suggesting a possible connection to the low contrast between blue and grey to their visual system. However, Streets et al. (2022) and our experiments both demonstrated that *H. trispinosa* can discriminate and learn blue from grey stimuli, indicating the results from Marshall et al. (1996) might be affected by other reasons, such as variations between different species or prolonged captivity of individuals in Marshall et al. (1996) that influenced the results. Nonetheless, our findings agree with Marshall et al. (1996) in highlighting blue as the most challenging stimulus among different colours. Why might this be?

#### (i) Potential low contrast between blue and grey stimuli

To investigate whether the contrast difference between colour and grey stimuli contributed to the stomatopod’s poor performance with the blue stimulus, we applied the multi-dichromatic processing scheme proposed by Marshall et al. (1996) to simulate the colour perception in stomatopods’ colour vision. It is suggested that the multi-dichromatic processing scheme is analogous to polarisation processing system, compares spectral instead of polarisation signals. How and Marshall (2014) developed a polarisation processing model designed for two-channel polarisation-sensitive system to measure the polarisation distance (contrast) between horizontal and vertical information of objects.

In stomatopods’ colour processing system, the two subsets (R1, 4, 5 and R 2, 3, 6, 7) of the R1-7 terminate at the separate layers in the lamina. The signals are presumed to be transmitted to the interneurons in the lamina through opponent connections. Accordingly, the activity profile (output) of the interneuron decoding a stimulus is represented by the equation:

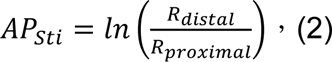

Rdistal and Rproximal are the quantum catches of the two subsets of receptors of R1-7 in a single row, with the receptor potential approximated by the natural log of the quantum catch. Rows 1-4 produce four separate activity profiles for one stimulus. The contrast difference (CD) of two stimuli to a row is then calculated by subtracting the activity profiles produced from decoding the two stimuli. For instance, the contrast difference of row 1 observing blue and grey stimuli is measured by the absolute difference between the activity profiles of row 1 in response to the blue and grey stimuli.

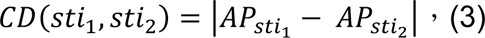

To determine whether the contrast difference of colour and grey stimuli to stomatopods’ colour vision aligns with the behavioural results, we measured the contrast differences between the colour and grey stimuli of each row under natural light and coloured illumination. Figure. 9A illustrates the contrast difference of the colour stimuli and 1.5 ND as an example. Additional comparison of CD values of colour and other grey stimuli can be found in Figure S1.

**Figure 9.**
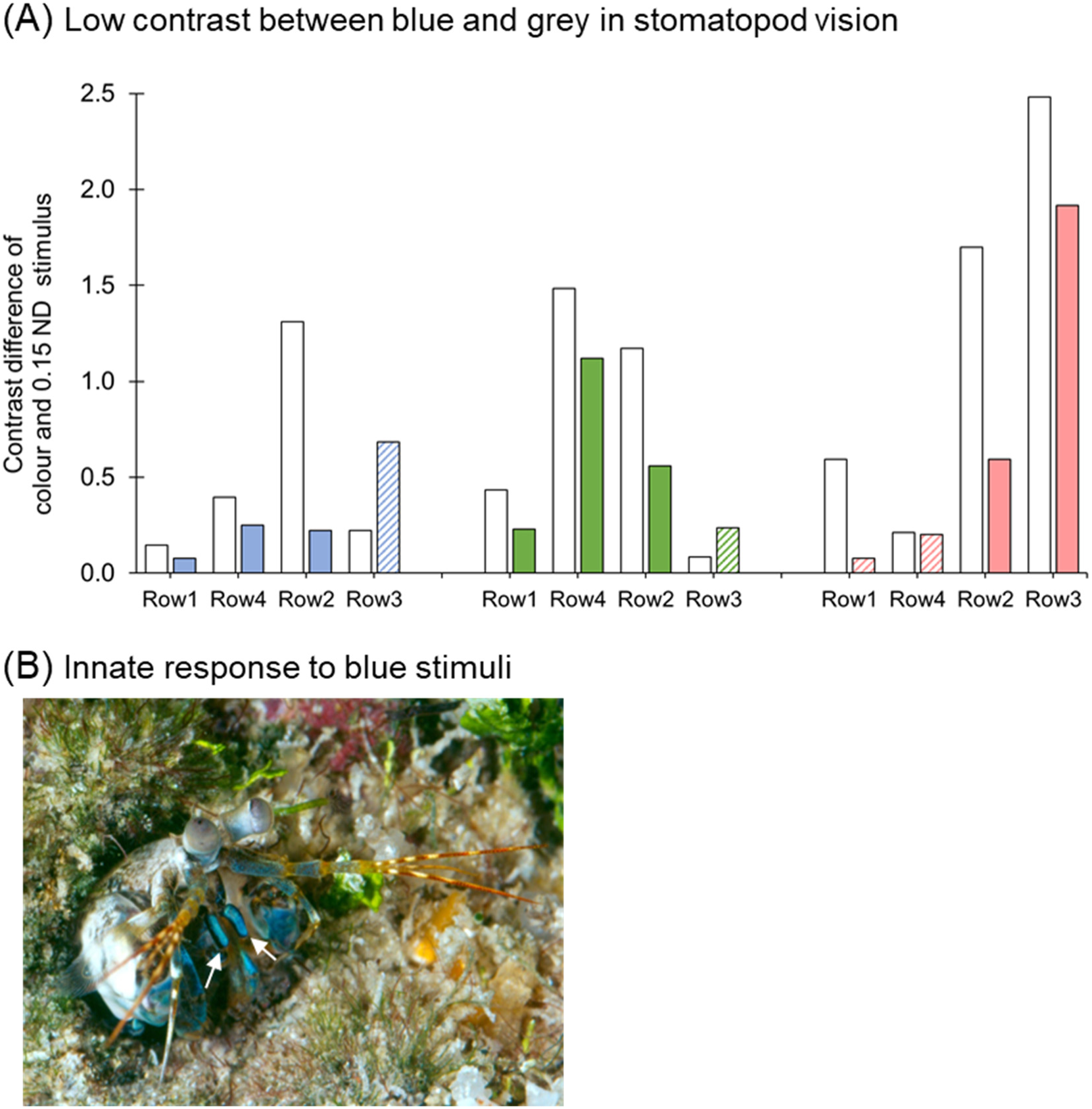
Potential factors contributing to behavioural variances of *H. trispinosa* with blue stimuli. (A) The figure illustrates the contrast difference (CD) between colour and 0.15 ND grey stimuli as determined by spectral rows 1-4 under both natural light (white bars) and coloured illuminations. The filled colour bar indicates the activated spectral row, while the coloured bars with lines represent the rows that were not activated under the corresponding coloured illumination. (B) Picture of *H. trispinosa*, showing blue reflection of the maxillipeds (arrows) and other body parts.

Generally, contrast differences between the colour and grey stimuli were higher in natural light than in coloured illuminations. Not surprisingly, colour stimuli appeared more distinct from greys to stomatopods under natural light than under coloured illumination, yet the contrast differences under coloured illuminations were still enough for green and red groups to distinguish the colours from grey stimuli.

However, while both green and red group had two rows with high CD for identifying colour under natural and coloured illumination, it appears that the blue group only had row 2 with higher CD for colour discrimination under natural light conditions. This finding supports the low trainability and cooperative rate of the blue group compared to the other two groups. Furthermore, although the differences between blue and grey stimuli were sufficient for identification under natural light, the extremely low contrast profile from all rows under blue illumination provides a convincing explanation for their failure to differentiate blue from grey stimuli under such conditions.

It is noteworthy that although row 3 seemed to present a relatively high CD under blue illumination, it contains only long-wavelength receptors, thus the signals perceived from row 3 under blue light should be extremely limited for effective signal processing. Due to the same reason, row 3 in the green tests and rows 1 and 4 in the red tests were most likely not effectively activated under the coloured illuminations; therefore, the CD values produced from these rows were presumed unhelpful for identifying colour under coloured illuminations.

All in all, this model supports the behavioural results (both here and in Marshall et al. (1996)), indicating that the discrepancy in the blue group’s behaviour is associated with the inadequate contrast between blue and grey stimuli under blue illumination in their visual system. Another significant finding is that the consistency between the results of the spectral processing model and the behavioural tests reinforces the idea of a multi-dichromatic processing system used for analysing spectral information in stomatopods’ colour vision.

#### (ii) Possible innate behaviours toward blue stimuli

An alternative explanation for the distinct performance in the blue group may be their innate response to blue stimuli, potentially influencing the choice tests.

Stomatopods inhabit environments rich in spectral and polarisation information, possessing a complex visual system evolved for fitness optimization. Certain signals are hardwired to elicit specific behaviours, aiding in conflict avoidance or enhancing reproductive success. Bok et al. (2018) discovered that *H. trispinosa* can discriminate different wavelengths of UV stimuli and exhibits an aversive response to UVB signals, possibly serving as a species recognition signal during encounters. Similarly, circular polarisation signals have been identified as burrow occupation cues, preventing territorial conflicts during burrow searches (Gagnon et al., 2015).

The maxillipeds of some stomatopods, especially in the genus Haptosquilla (How et al., 2014), commonly reflect blue signals, sometimes with strong polarisation (Fig. 9B). Chiou et al. (2011) revealed that the blue-polarized reflection from *H. trispinosa* served as mating signals. Female *H. trispinosa* showed a preference for mating with males possessing blue-polarized maxillipeds, while it took longer for females to accept control males, whose maxillipeds were covered with black paint. These findings imply that *H. trispinosa* may associate blue cues strongly with species recognition, perhaps making it a unfavored signal for food choice.

In Streets et al. (2022), naive tests revealed intriguing behaviours in *H. trispinosa* regarding their responses to various stimuli. While they didn’t exhibit aversion to saturated blue stimuli compared to saturated red, orange, and green ones, they did display a preference for low-saturated blue when presented with both saturated and unsaturated blue stimuli. Similarly, in naive tests comparing saturated blue with grey, stomatopods favoured grey over saturated blue. These findings indicated that saturated blue stimuli may elicit innate avoidance behaviours, particularly when presented alongside stimuli with low saturation levels, such as unsaturated blue or grey stimuli.

While the interplay of polarisation, colour and brightness information complicates determining whether the reflectance of *H. trispinosa*’s maxillipeds resembles unsaturated or saturated blue in their visual system. Nonetheless, given their evident avoidance toward saturated blue stimuli, it cannot be ruled out that their innate behaviours may contribute to the learning challenges and poor performance observed in the blue group.

### (3) Information processing in stomatopods colour vision

In Thoen et al. (2014), it was proposed that the multi-dichromatic processing system compares adjacent spectral sensitivities would enable a spectral discrimination curve with overall Δλ value below 5 nm in their colour vision (Fig. 2D) (Thoen et al., 2014). However, the fluctuating discrimination in behavioural observations indicated a tetrachromatic-like system with low resolution, exhibiting over three times higher Δλ value than expected (Fig. 10A). This suggests that stomatopods may possess tetrachromatic vision but without a fine or thorough comparison as proposed in the visual model.

**Figure 10.**
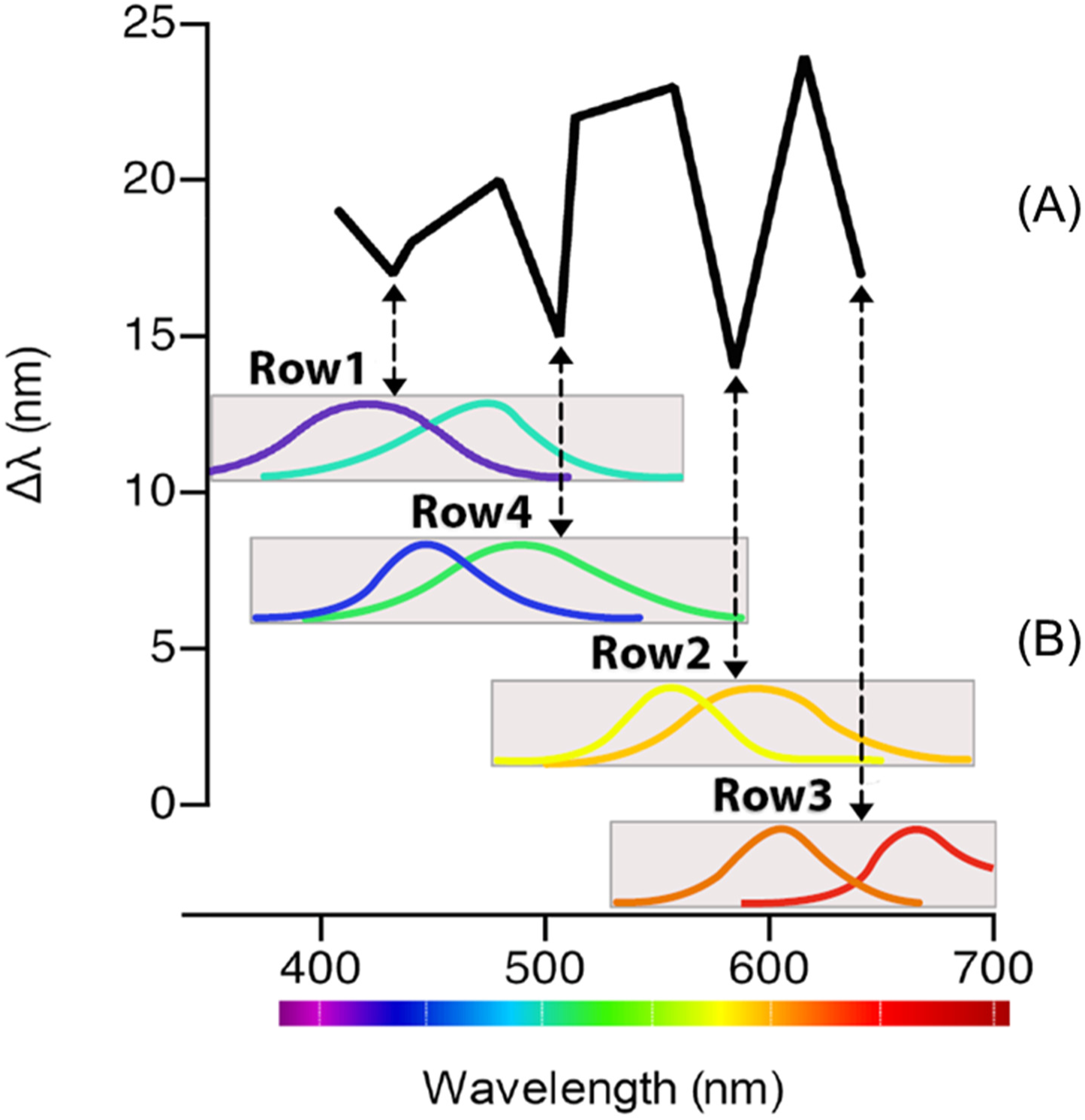
Spectral discrimination curve alignment with multi-dichromatic processing system in *H. trispinosa*. The spectral discrimination curve, as observed in *H. trispinosa* (Thoen et al., 2014), reveals four distinct dips indicative of enhanced spectral resolution in the animals. These four dips align closely with the four spectral window samples represented by spectral rows 1-4, supporting multi-dichromatic processing. Additionally, the lack of fine spectral resolution suggests that the outputs from the four spectral windows may undergo binning processing at the downstream visual pathway.

Our findings support a hybrid system in stomatopods’ colour vision, combining opponent and binning mechanisms. It is hypothesised this system compares spectral information within each spectral row (rows 1-4), establishing four “processed spectral sensitivities” that form a tetrachromatic pattern (Fig. 10B). Subsequently, these outputs may be relayed downstream through a binning system without complex comparison, leading to coarse colour resolution.

This hypothesis gains support from the neuronal constructions of their midband processing pathway along the optic lobes, proposing a combined system where opponent processing governs the first optic lobe (the lamina), while binning processing dominates in the downstream optic lobes (the medulla and lobula) (Thoen et al., 2017a; Thoen et al., 2018). The close resemblance of the arrangement of the lamina of rows 1-4 and the polarisation system in crustaceans indicates the use of an opponent processing scheme for decoding spectral information. Meanwhile, the organisation of the medulla and lobula in stomatopods appears to align with the concept of binning system. Unlike insects, where medulla is the first stage the spectral information is pooled and compared (Paulk et al., 2008; Paulk et al., 2009; Schnaitmann et al., 2018; Schnaitmann et al., 2020), stomatopods may exhibit spectral opponency in the lamina, with limited spectral comparison the medulla. The midband lamina terminals project to a the midband track comprised of four chromatic visual sampling units in the medulla. The midband track cuts through the stratified organisation of the medulla and propagates to deep lobula, forming a distinguished pathway from the hemisphere representation (Thoen et al., 2018). Such a direct pathway from the lamina to deep levels of lobula implies that a lack of complex comparison between different spectral rows, supporting the idea of binning processing in these neuropils. Further work, especially electrophysiology study, is required to gain a better understanding of the processing mechanism along their spectral processing pathway.

Another complex colour vision system, butterflies *Papilio xuthus*, possess up to eight classes of spectrally distinct photoreceptors, making it the only other animal with comparable retinal complexity to stomatopods (Arikawa et al., 1987; Arikawa, 2003; Koshitaka et al., 2008, Marshall and Arikawa 2014). Previous studies discovered a special early spectral processing in the retinal level of butterflies (Chen et al., 2013; Chen et al., 2019; Chen et al., 2020), with the spectral opponent processing in butterflies take place as early as the receptor terminal. The spectral discrimination ability tests have shown that butterflies are tetrachromatic and have the smallest known Δλ values (approximately 1 nm) among animals (Koshitaka et al., 2008). It is suggested that the early stage of visual processing in butterflies might help to streamline the signals and lead to more effective processing of information in their visual system. Likewise, with even more spectral channels plus polarisation vision in their visual system, stomatopods are expected to also face the challenge of handling complex incoming data from their retina. Such pre-processing and organization of spectral signals in the early stage of processing, like those of stomatopods and butterflies, may help to minimise the computational load in the deeper and more complex processing centre.

Our finding demonstrated that stomatopods probably possess four opponent channels in their colour vision, which finds support from the proposal that colour vision evolved to possess spectral opponency to eliminate flicker in shallow water environments (Maximov, 2000). Such an ability is particularly advantageous for animals like stomatopods as many live in very shallow-water environments, where the reduction of flicker may enhance efficiency in prey and threat detection.

Additionally, the multi-dichromatic colour vision has been previously hypothesised to be and adaptation to enhance colour constancy in the aquatic environment in stomatopods’ colour vision Osorio et al. (1997). By simulating stomatopods’ colour vision multiple opponent spectral rows, Osorio et al. (1997) suggested the multi-dichromatic system enables them to decode information within a narrow spectral window. This process potentially allows stomatopods to obtain consistent colour perception under varying lighting conditions, hinting colour constancy in their colour vision. Future studies on stomatopods’ colour constancy are anticipated to provide important insights into their unique colour vision system and its adaptive significance in their ecological niche.

## Acknowledgements

We are grateful for extensive field trip assistance from members of the Marshall laboratory (notably Dr. Fabio Cortesi, Dr. Wen-Sung Chung and Abi Shaughnessy) and The Australian Museum’s Lizard Island Research Station. Thank Dr. Tsyr-Huei Chiou for providing the stomatopod eye picture. Special thanks to Dr. Amy Streets for sharing her protocols and expertise in behavioural procedures and Dr. Lily Fogg for assisting Experiment 2.

## Author contributions

C.W.J.W, and J.M. conceived and designed this study. C.W.J.W conducted the experiments, handled the data curation and performed the data analysis and interpretation. Both authors validated the results of the data analysis and visual model. The manuscript was initially drafted by C.W.J.W and edited by J.M. and C.W.J.W. J.M. provided project supervision, acquired research funding and managed project administration.

## References

Arikawa, K. (2003). Spectral organization of the eye of a butterfly, Papilio. J. Comp. Physiol. A 189, 791–800.

Arikawa, K. and Stavenga, D. G. (2014). Insect Photopigments: Photoreceptor Spectral Sensitivities and Visual Adaptations. In Evolution of Visual and Non-visual Pigments (ed. Hunt, D. M.), Hankins, M. W.), Collin, S. P.), and Marshall, N. J.), pp. 137–162. Boston, MA: Springer US.

Arikawa, K., Inokuma, K. and Eguchi, E. (1987). Pentachromatic visual system in a butterfly. Naturwissenschaften 74, 297–298.

Barlow, H. B. (1982). What causes trichromacy? A theoretical analysis using comb-filtered spectra. Vision Res. 22, 635–643.

Bok, M. J., Roberts, N. W. and Cronin, T. W. (2018). Behavioural evidence for polychromatic ultraviolet sensitivity in mantis shrimp. Proc. R. Soc. B Biol. Sci. 285, 20181384.

Chen, P.-J., Arikawa, K. and Yang, E.-C. (2013). Diversity of the Photoreceptors and Spectral Opponency in the Compound Eye of the Golden Birdwing, Troides aeacus formosanus. PLOS ONE 8, e62240.

Chen, P.-J., Matsushita, A., Wakakuwa, M. and Arikawa, K. (2019). Immunolocalization suggests a role of the histamine-gated chloride channel PxHCLB in spectral opponent processing in butterfly photoreceptors. J. Comp. Neurol. 527, 753–766.

Chen, P.-J., Belušič, G. and Arikawa, K. (2020). Chromatic information processing in the first optic ganglion of the butterfly Papilio xuthus. J. Comp. Physiol. A 206, 199–216.

Cheroske, A. G. and Cronin, T. W. (2005). Variation in Stomatopod *(Gonodactylus smithii)* Color Signal Design Associated with Organismal Condition and Depth. Brain. Behav. Evol. 66, 99–113.

Cheroske, A. G., Cronin, T. W. and Caldwell, R. L. (2003). Adaptive color vision in Pullosquilla litoralis (Stomatopoda,Lysiosquilloidea) associated with spectral and intensity changes in light environment. J. Exp. Biol. 206, 373–379.

Chiou, T.-H., Kleinlogel, S., Cronin, T., Caldwell, R., Loeffler, B., Siddiqi, A., Goldizen, A. and Marshall, J. (2008). Circular Polarization Vision in a Stomatopod Crustacean. Curr. Biol. 18, 429–434.

Chiou, T.-H., Marshall, N. J., Caldwell, R. L. and Cronin, T. W. (2011). Changes in light-reflecting properties of signalling appendages alter mate choice behaviour in a stomatopod crustacean Haptosquilla trispinosa. Mar. Freshw. Behav. Physiol. 44, 1–11.

Cronin, T. W. and Caldwell, R. (2002). Tuning of photoreceptor function in three mantis shrimp species that inhabit a range of depths. II. Filter pigments. J. Comp. Physiol. [A] 188, 187–197.

Cronin, T. W. and Marshall, N. J. (1989a). A retina with at least ten spectral types of photoreceptors in a mantis shrimp. Nature 339, 137–140.

Cronin, T. W. and Marshall, N. J. (1989b). Multiple spectral classes of photoreceptors in the retinas of gonodactyloid stomatopod crustaceans. J. Comp. Physiol. A 166,.

Cronin, T. W. and Marshall, N. J. (2001). Parallel Processing and Image Analysis in the Eyes of Mantis Shrimps. Biol. Bull. 200, 177–183.

Cronin, T. W., Caldwell, R. L. and Marshall, N. J. (2001). Tunable colour vision in a mantis shrimp. Nature 411, 547–548.

Cronin, T. W., Caldwell, R. L. and Erdmann, M. V. (2002). Tuning of photoreceptor function in three mantis shrimp species that inhabit a range of depths. I. Visual pigments. J. Comp. Physiol. A 188, 179–186.

Daly, I. M., How, M. J., Partridge, J. C., Temple, S. E., Marshall, N. J., Cronin, T. W. and Roberts, N. W. (2016). Dynamic polarization vision in mantis shrimps. Nat. Commun. 7, 12140.

De Valois, R. L. and Jacobs, G. H. (1968). Primate Color Vision. Science 162, 533– 540.

Frisch, K. von (1914). Der farbensinn und Formensinn der Biene.

Gagnon, Y. L., Templin, R. M., How, M. J. and Marshall, N. J. (2015). Circularly Polarized Light as a Communication Signal in Mantis Shrimps. Curr. Biol. 25, 3074–3078.

Glantz, R. M. and Miller, C. S. (2002). Signal Processing in the Crayfish Optic Lobe: Contrast, Motion and Polarization Vision. In The Crustacean Nervous System (ed. Wiese, K.), pp. 486–498. Berlin, Heidelberg: Springer Berlin Heidelberg.

How, M. J. and Marshall, N. J. (2014). Polarization distance: a framework for modelling object detection by polarization vision systems. Proc. R. Soc. B Biol. Sci. 281, 20131632.

How, M. J., Porter, M. L., Radford, A. N., Feller, K. D., Temple, S. E., Caldwell, R. L., Marshall, N. J., Cronin, T. W. and Roberts, N. W. (2014). Out of the blue: the evolution of horizontally polarized signals in *Haptosquilla* (Crustacea, Stomatopoda, Protosquillidae). J. Exp. Biol. jeb.107581.

Kelber, A., Vorobyev, M. and Osorio, D. (2003). Animal colour vision – behavioural tests and physiological concepts. Biol. Rev. Camb. Philos. Soc. 78, 81–118.

Kleinlogel, S. and Marshall, N. J. (2005). Photoreceptor projection and termination pattern in the lamina of gonodactyloid stomatopods (mantis shrimp). Cell Tissue Res. 321, 273–284.

Kleinlogel, S., Marshall, N. J., Horwood, J. M. and Land, M. F. (2003). Neuroarchitecture of the color and polarization vision system of the Stomatopod haptosquilla. J. Comp. Neurol. 467, 326–342.

Koshitaka, H., Kinoshita, M., Vorobyev, M. and Arikawa, K. (2008). Tetrachromacy in a butterfly that has eight varieties of spectral receptors. Proc. R. Soc. B Biol. Sci. 275, 947–954.

Maloney, L. T. (1986). Evaluation of linear models of surface spectral reflectance with small numbers of parameters. J. Opt. Soc. Am. A 3, 1673.

Manning, R. B. (1977). A monograph of the West African stomatopod Crustacea. Copenhagen: Scandinavian Science Press.

Marshall, N. J. (1988). A unique colour and polarization vision system in mantis shrimps. Nature 333, 557–560.

Marshall, N. J. and Arikawa, K. (2014). Unconventional colour vision. Curr. Biol. 24, R1150–R1154.

Marshall, N. J. and Land, M. F. (1993). Some optical features of the eyes of stomatopods: II. Ommatidial design, sensitivity and habitat. J. Comp. Physiol. A 173, 583–594.

Marshall, N. J. and Oberwinkler, J. (1999). The colourful world of the mantis shrimp. Nature 401, 873–874.

Marshall, N. J., Land, M. F., King, C. A. and Cronin, T. W. (1991). The compound eyes of mantis shrimps (Crustacea, Hoplocarida, Stomatopoda). I. Compound eye structure: the detection of polarized light. Philos. Trans. R. Soc. Lond. B. Biol. Sci. 334, 33–56.

Marshall, N. J.., Jones, J. and Cronin, T. W. (1996). Behavioural evidence for colour vision in stomatopod crustaceans. J. Comp. Physiol. A 179, 473–481.

Marshall, N. J., Cronin, T. W. and Kleinlogel, S. (2007). Stomatopod eye structure and function: A review. Arthropod Struct. Dev. 36, 420–448.

Maximov, V. V. (2000). Environmental factors which may have led to the appearance of colour vision. Philos. Trans. R. Soc. B Biol. Sci. 355, 1239–1242.

Neumeyer, C. (1986). Wavelength discrimination in the goldfish. J. Comp. Physiol. A 158, 203–213.

Osorio, D., Marshall, N. J. and Cronin, T. W. (1997). Stomatopod photoreceptor spectral tuning as an adaptation for colour constancy in water. Vision Res. 37, 3299–3309.

Paulk, A. C., Phillips-Portillo, J., Dacks, A. M., Fellous, J.-M. and Gronenberg, W. (2008). The Processing of Color, Motion, and Stimulus Timing Are Anatomically Segregated in the Bumblebee Brain. J. Neurosci. 28, 6319–6332.

Paulk, A. C., Dacks, A. M. and Gronenberg, W. (2009). Color processing in the medulla of the bumblebee (Apidae: Bombus impatiens). J. Comp. Neurol. 513, 441–456.

Porter, M. L., Zhang, Y., Desai, S., Caldwell, R. L. and Cronin, T. W. (2010). Evolution of anatomical and physiological specialization in the compound eyes of stomatopod crustaceans. J. Exp. Biol. 213, 3473–3486.

Schnaitmann, C., Haikala, V., Abraham, E., Oberhauser, V., Thestrup, T., Griesbeck, O. and Reiff, D. F. (2018). Color Processing in the Early Visual System of Drosophila. Cell 172, 318–330.e18.

Schnaitmann, C., Pagni, M. and Reiff, D. F. (2020). Color vision in insects: insights from Drosophila. J. Comp. Physiol. A 206, 183–198.

Strausfeld, N. and Nassel, D. (1981). Neuroarchitectures serving compound eyes of Crustacea and insects. In Handbook of sensory physiology, pp. 34–41. Springer.

Streets, A., England, H. and Marshall, J. (2022). Colour vision in stomatopod crustaceans: more questions than answers. J. Exp. Biol. 225, jeb243699.

Templin, R. (2017). Circular polarization vision in stomatopod crustaceans.

Thoen, H. H., How, M. J., Chiou, T.-H. and Marshall, N. J. (2014). A Different Form of Color Vision in Mantis Shrimp. Science 343, 411–413.

Thoen, H. H., Strausfeld, N. J. and Marshall, N. J. (2017a). Neural organization of afferent pathways from the stomatopod compound eye. J. Comp. Neurol. 525, 3010–3030.

Thoen, H. H., Chiou, T.-H. and Marshall, N. J. (2017b). Intracellular Recordings of Spectral Sensitivities in Stomatopods: a Comparison across Species. Integr. Comp. Biol. 57, 1117–1129.

Thoen, H. H., Sayre, M. E., Marshall, N. J. and Strausfeld, N. J. (2018). Representation of the stomatopod’s retinal midband in the optic lobes: Putative neural substrates for integrating chromatic, achromatic and polarization information. J. Comp. Neurol. 526, 1148–1165.

von Helversen, O. (1972). Zur spektralen Unterschiedsempfindlichkeit der Honigbiene. J. Comp. Physiol. 80, 439–472.

